# Swarming bacteria respond to increasing barriers to motility by increasing cell length and modifying colony structure

**DOI:** 10.1101/398321

**Authors:** Kristin Little, Jacob L. Austerman, Jenny Zheng, Karine A. Gibbs

## Abstract

Organisms can alter morphology and behaviors in response to environmental stimuli such as mechanical forces exerted by surface conditions. The bacterium *Proteus mirabilis* responds to surface-based growth by enhancing cell length and degree of cell-cell interactions. Cells grow as approximately 2-micrometer rigid rods and independently swim in liquid. By contrast on hard agar surfaces, cells elongate up to 40-fold into snake-like cells that move as a collective group across the surface. Here we have elucidated that individual cell size and degree of cell-cell interactions increased across a continuous gradient that correlates with increasing agar density. We further demonstrate that interactions between the lipopolysaccharide (LPS) component of the outer membrane and the immediate local environment modified these responses by reducing agar-associated barriers to motility. Loss of LPS structures corresponded with increased cell elongation on any given surface. These micrometer-scale changes to cell shape and collective interactions corresponded with centimeter-scale changes in the overall visible structure of the swarm colony. It is well-appreciated in eukaryotes that mechanical forces impact cell shape and migration. Here we propose that bacteria can also dynamically respond to the mechanical forces of surface conditions by altering cell shape, individual motility, and collective migration.

## Introduction

Eukaryotic and prokaryotic cells undergo alterations, including changes in cell morphology and collective behaviors, in response to interactions with the physical environment. For example, robust swarmers such as the human pathogens *Proteus mirabilis* and *Vibrio parahaemolyticus* (1) can swim as short, rigid rod-shaped cells in liquid and through low-density (0.3%) agar. On low-wetness and highdensity agar (0.75% to 2.5%), these bacteria elongate dramatically and move as a collective group on top of the surface. By comparison, the swarm motility of temperate swarmers such as *Escherichia coli* or *Salmonella enterica* is generally restricted to low-density agar (< 0.7%) or high wetness Eiken agar (reviewed in (2, 3)). For all of the aforementioned bacteria, flagella power the motility in liquid and on surfaces. Furthermore, a transition from swimming to swarming coincides with a series of large-scale transcriptional changes triggered by contact between flagella and a surface (Figure 1) (4–12).

**Fig. 1.**
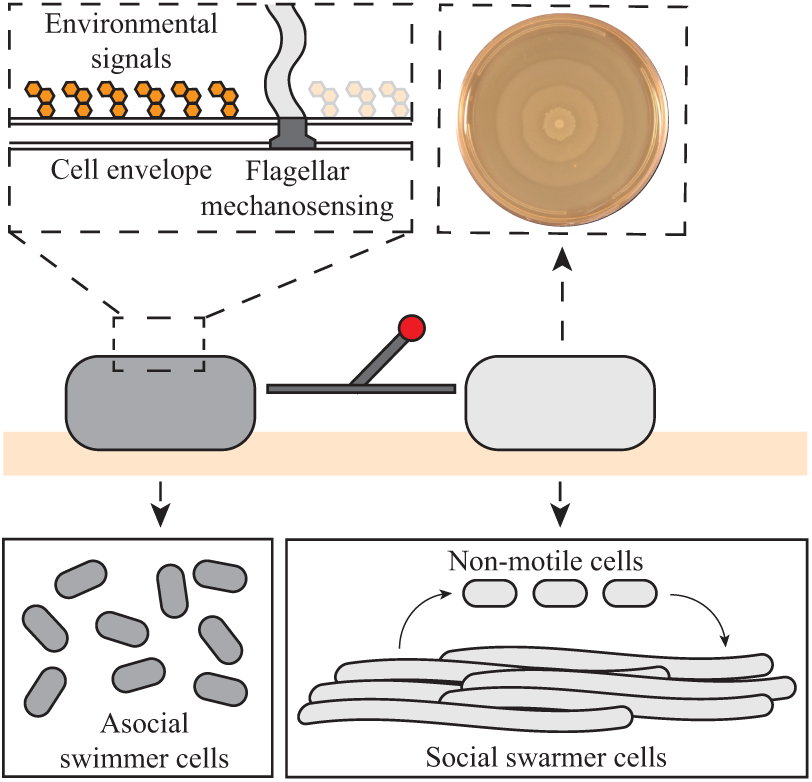
Current model for surface adaptation. *P. mirabilis* cells can adopt at least two morphologically and behaviorally distinct lifestyles based on environmental conditions. In liquid, cells exist as short, non-social swimming cells. Upon contact with a surface, cells switch into the swarming lifestyle. Swarmer cells dramatically elongate, form multicellular groups, and move across surfaces to form terraced colonies. After a period of motility, cells divide into short, non-motile cells that remain behaviorally and transcriptionally distinct from swimming cells.

Here we utilize *P. mirabilis* as a tractable model for exploring how morphology and cell-cell interactions respond to the surface. *P. mirabilis* cells are approximately 2-µm long, rigid, and rod-shaped in liquid. Such cells swim independently, resulting in a visible uniform haze to the structure of the swim colony. Upon contact with a hard surface (Figure 1), these rigid, rod-shaped cells can elongate up to 40-fold into a flexible, snake-like, hyper-flagellated swarmer cell (13, 14) that has a distinctive gene expression profile (15). *P. mirabilis* swarmer cells are thought to bundle their flagella to facilitate cooperative swarm motility on hard agar (16). Rafting appears to be important for motility, as cells that escape a raft exhibit reduced motility. After a period of motility, swarmer cells divide into rigid, 2-µm long, non-swarm-motile cells (Figure 1). Iterative cycles of elongation, motility, and division result in a visible stereotyped bullseye-patterned swarm colony that can occupy several centimeters. Each wide-diameter ring, herein referred to as a terrace, is thought to reflect the expanding migratory period of elongated cells. The darker perimeter circles reflect the division period and is occupied by short, non-swarm-motile cells.

One critical factor for this swarm developmental cycle is lipopolysaccharide (LPS) biosynthesis and structure (17, 18). LPS, which comprises the outer leaflet of the outer membrane of Gram-negative bacteria, is also a critical requirement for surface-based motility in several additional bacterial species such as *E. coli* (19), *Myxococcus xanthus* (20, 21), and *S. enterica* (22, 23). Disruption of LPS biosynthesis genes, including the O-antigen ligase *waaL* (18, 24) and the sugar-modifying enzyme *ugd* (17, 25), has been shown to inhibit swarmer cell development through activation of cell envelope stress-associated pathways in *P. mirabilis*. In addition, LPS is hypothesized to physically promote swarm motility by acting as a wetting agent shared between cells (26–30). Specifically, LPS is suggested to serve as an osmolarity agent to draw moisture from an agar gel environment and facilitate swarming in *S. enterica* (22, 23). A critical aspect of LPS swarm-related functions remains unclear: whether LPS acts locally to each cell or is shared across multiple cells to potentially enhance the motility of neighboring cells. In one model, LPS is hypothesized to detach from cells and serve as a shared good to enhance wetness, thereby increasing the ability of the entire colony to expand outwards (31). Indeed, the roles of LPS in swarm motility and cell development are unreconciled.

We addressed this intersection of development and motility using *P. mirabilis* as it is able to adopt morphologically, physiologically, and behaviorally distinct lifestyles in liquid and surface environments. Here we provide original evidence that surface-grown *P. mirabilis* cells actually adopt a gradient of cell lengths and of cell-cell interactions in response to variations in agar concentrations. We further demonstrate that LPS plays a critical role in the swarming environment of *P. mirabilis*, perhaps by aiding cells to migrate on high-density agar. We found that this LPS-mediated activity occurs local to individual cells. We determined that LPS function correlates with surface-associated changes in cell morphology: cells with defective LPS structure elongated more extensively on surfaces than wildtype. Moreover, we found that microscopic changes in cell morphology and cellcell interactions correlated with centimeter-scale changes to colony structure for both wildtype and LPS-defective cells. We propose that cells increase cell shape and degree of cellcell interactions as a dynamic response to barriers to swarm motility and that LPS serves to lower such barriers by altering a cell’s local interactions with the surface. This research begins to establish that micron-scale cell morphology, cell-cell interactions, and localized cell-environment interactions together influence the formation of centimeter-scale structures.

## Results

### Swarmer cell size, cell-cell interactions, and colony structure shift in response to agar concentration

We set out to examine how *P. mirabilis* populations of strain BB2000 acclimate to surface-grown conditions. We chose to alter a single surface variable— agar percentage— to determine whether the visible structure of swarm colonies, cell morphology, or cell-cell interactions responded to changes in surface properties. *P. mirabilis* swarms are typically grown on nutrient-rich medium, e.g., Lennox lysogeny (LB) or CM55 containing 1.5% agar. Increasing agar concentration up to 2.5% is known to narrow swarm colony terraces and beyond 2.5% agar, to inhibit swarm colony expansion (28).

Using LB medium, we altered the agar concentrations, ranging from 0.75%, on top of which cells move, to 4%, on which cells form isolated colonies on top of the surface. Wild-type strain BB2000 cells were inoculated at a single point on LB plates followed by growth at 37°C overnight. Populations grew to form unstructured thin films on top of the 0.75% agar plates (Figure 2A). This visible structure is distinct from the stereotypical haze of a swimming population grown in 0.3% agar (32). Terraces were visible between 1% agar and 2.5% agar and narrowed in width as agar concentration increased (Figure 2A). On 1 and 1.25% agar, populations formed what we termed, “semi-structured swarm colonies,” in which visible terraces were broad and poorly defined (Figure 2A). On 1.5%, 1.75%, and 2% agar, populations formed distinctly terraced swarm colonies with terrace width inversely correlated with agar concentration (Figure 2A, Supplemental Figure SF1). On 2.5% agar, the terrace rings became indistinct, lending the colony a mucoid appearance (Figure 2A). Populations did not expand beyond the inoculation point when the agar concentration was 4% (Figure 2A). Thus, the formation of visible terraced structures, as well as narrowing of terrace width, emerged at 1% agar and increased in frequency as the agar concentration increased.

**Fig. 2.**
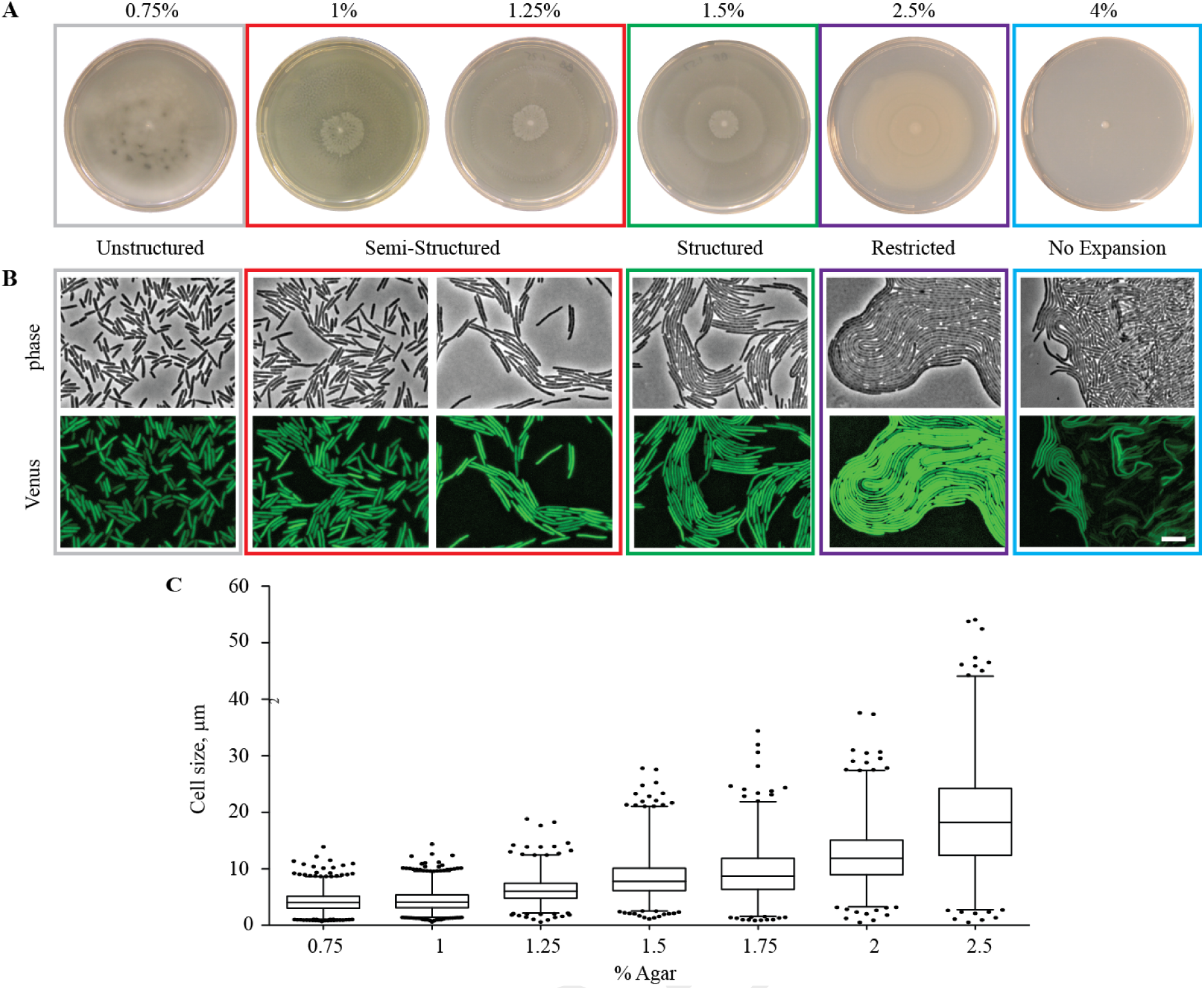
Swarmer cell length and colony structure scale with agar concentration. A. Representative images of populations inoculated onto LB medium containing 0.75 4.0% agar and grown overnight at 37°C on a 10-cm diameter petri dish. Colony structure was categorized as unstructured, semi-structured, structured, and restricted based on comparison of terrace definition and width relative to wild-type populations grown on 1.5% agar medium. B. Representative images of cells encoding a chromosomal Venus reporter for *fliA* expression. Cells were inoculated onto 1-mm thick LB medium pads containing indicated concentrations agar, grown at 37°C, and imaged using epifluorescence microscopy. Images include images in phase (top) and fluorescence (bottom). Fluorescence images were subjected to background subtraction. The corresponding macroscopic swarm colony structure characteristics as described in (A) are indicated above. Scale bars, 10 µm. C. Swarmer cell size was measured on the denoted media conditions. Wild-type cells were co-mixed in a 10:1 ratio with wild-type cells carrying the *fliA* reporter strain. Data are shown in box and whisker plots; the box denotes median, 25th, and 75th percentiles. Whiskers boundary 1st and 99th percentiles. Individual data points represent values beyond the 1 to 99 percentile range. The numbers of measured cells for each agar percentage, sequentially from 0.75% to 2.5% agar, are as follows: 2446, 2894, 1419, 1502, 1262, 1263, and 913.

We considered whether shifts toward structured colonies and narrower swarm rings with increasing agar concentration resulted from changes in single-cell morphology or behavior. We used epifluorescence microscopy to examine individual cells and followed the initiation of the swarm developmental cycle by monitoring a Venus reporter for *fliA* expression; this reporter was integrated onto the chromosome. The *fliA* gene encodes sigma 28 factor and is positively regulated by FlhD_4_C_2_ (33). This reporter strain has no discernible differences in swarm behavior as compared to the unlabeled parent strain. Briefly, when cells elongate into flexible, motile swarmer cells, fluorescence associated with the *fliA* reporter is highly exhibited (32). Such swarming cells often migrate across the surface in multicellular packs termed rafts (Figure 2B). The reporter strain was grown on LB agar conditions ranging from 0.75% to 4% agar. We observed that the length of each swarmer cell and the numbers of cells per group increased as the agar concentration increased (Figure 2B). Cells did not consistently align into groups of cells when grown on 0.75% to 1% agar (Figure 2B). However, groups of cells aligned with one another and the number of cells within each raft increased as the agar concentration increased from 1.25% to 2.5% (Figure 2B). Corresponding movies can be found on the K. A. Gibbs’ Research Group’s YouTube channel.

To quantify swarmer cell size, we mixed unlabeled wild-type cells with cells carrying the fluorescent *fliA* reporter at a 10:1 ratio, allowed populations to initiate swarm motility, and imaged across the swarm front. The area of the fluorescent signal associated with each individual swarmer cell within larger clusters was measured. Cells grown on 0.75% to 1% agar remained relatively short, with median cell areas of 4.0 and 4.1 µm^2^ respectively (Figure 2C). Cells became increasingly elongated as agar concentration increased from 1.25% 2.5% agar. Cells on the swarm-restrictive 2.5% agar were notably longer that cells grown on 1.5% agar, with a median cell area of 18.2 µm^2^ versus 7.8 µm^2^, respectively. The distribution of cell sizes increased as agar concentration increased. Therefore, we concluded that swarmer cells were becoming increasingly heterogeneous in length and more likely to form cell-cell contacts. The visible structure of the swarm colony also changed with increasing agar concentration; the terrace width decreased as agar concentration increased. We propose that increasing individual cell length, broadening of cell length distributions, and increasing the extent of cell-cell interactions correlated with the narrowing of swarm colony terraces.

### Perturbing LPS structures uncouples swarmer cell elongation from swarm colony expansion

We observed that an increase in agar concentration could functionally uncouple swarmer cell elongation from the outward-expanding motility of the total colony. Specifically, elongated wild-type cells were apparent on the periphery of colonies grown on 4% agar (Figure 2B) even though swarm colony expansion was not visible (Figure 2A). We reasoned that previously identified non-swarming mutant strains might actually be capable of elongating into swarmer cells. Such hypothetical mutant strains would fail to engage in swarm colony expansion on standard medium but would differentiate into swarmer cells. Such mutant strains could provide a factor(s) necessary for initiating swarm colony expansion after elongation.

We therefore analyzed non-swarming mutant strains that arose from a library of approximately 13,000 mutant strains. The library was generated from two independent transposon mutagenesis screens in *P. mirabilis* strain BB2000 (this study and (34)). A secondary screen was performed on each non-swarming mutant strain to look for elongated swarmer cells on swarm-permissive CM55 agar (nutrient-rich 1.5% agar). Two mutant strains (KEL21 and KEL22) were identified; each failed to engage in swarm colony expansion and elongated into swarmer cells on CM55 agar (Supplemental Figure SF2). The disruptions in strains KEL21 and KEL22 were mapped using a modified inverse Polymerase Chain Reaction protocol followed by Sanger sequencing. Strain KEL21 contains a disruption in the gene BB2000_3203 (Supplemental Figure SF2), which encodes a putative O-antigen acetylase. Strain KEL22 contains a disruption in the gene BB2000_3208 (Supplemental Figure SF2), which encodes a predicted NAD-dependent epimerase/dehydratase. Both genes are contained within a single locus for genes of an LPS biosynthesis pathway (Supplemental Figure SF2), which varies in the number and composition of genes among *P. mirabilis* strains (35).

We confirmed the predicted disruption of the LPS structures by extracting LPS from each mutant strain and the wild-type parent strain. The resulting fractions were analyzed using SDS-PAGE and a modified silver stain protocol (36). Comparably less LPS staining and nearly complete loss of the O-antigen ladder was repeatedly apparent in samples extracted from the strains KEL21 and KEL22 as compared to wild-type strain (Figure 3). Therefore, LPS production was disrupted in each mutant strain; the strains are likely deficient in the production of O-antigen. To evaluate the effects of these LPS deficiencies on cell envelope integrity, we examined the susceptibility of all strains to polymyxin B on swarm-permissive surfaces using antibacterial halo assays. In *P. mirabilis*, loss of LPS increases a cell’s sensitivity to the antimicrobial peptide polymyxin B (37, 38). Wildtype was resistant to a completely saturated solution of 50 mg/mL polymyxin B as expected. By contrast, both mutant strains were inhibited on surfaces by minimally 500 µg/mL. The mutant strains are approximately 100-fold more sensitive to polymyxin B than wildtype. We concluded that strains KEL21 and KEL22 each contained deficiencies in LPS structures.

**Fig. 3.**
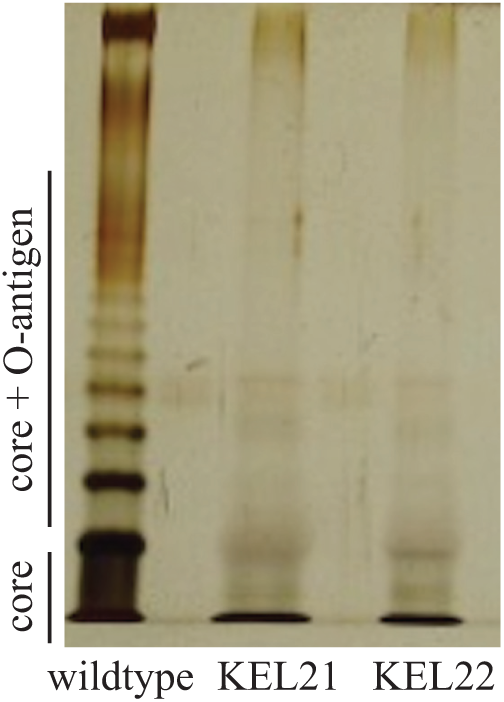
LPS-defective populations show disrupted LPS structures. LPS was independently extracted from surface-grown populations of strains wild type KEL21 and KEL22. These extracts were run on an SDS-PAGE gel and visualized with silver stain. For increased visualization, 5-fold (KEL21) and 2.5-fold (KEL22) excess of LPS from mutant strains was loaded relative to wildtype. Predicted sizes for lipid A core and O-antigen ladder are labeled on left. Representative image of three biological replicates.

### LPS-defective cells respond in cell shape and colony expansion at a lower range of agar concentrations than wildtype

Given these data, LPS structure might be a factor for initiating swarm colony expansion after elongation. Strains KEL21 and KEL22 were therefore subjected to analysis of swarm colony structure and single-cell morphology similar to those used for wildtype (Figure 2). For single-cell analysis, we constructed derived strains of KEL21 and KEL22 that encoded a chromosomal reporter for *fliA* expression; these cells were observed in clonal swarms and in mixed swarms with wild-type for size measurements (Figure 4). We evaluated a range of LB agar concentrations from 0.75% to 1.5%. At 0.75% agar, these populations covered the surface of the plate without apparent terrace structures (Figure 4). KEL21 and KEL22 cells on 0.75% agar were visibly shorter and more rigid. They had a median cell area of 5.4 and 5.7 µm^2^, respectively (Figure 4), which is comparable in size to wildtype cells grown on 1.25% agar (Figure 4). By contrast to wildtype, we observed that swarm colonies of both mutant strains exhibited visible terraces on agar concentrations of 1% and 1.25% (Figure 4). Flexible, elongated cells that aligned into clusters of cells were readily visible within these swarm colonies with a median cell area of approximately 10 µm^2^ and 13 µm^2^, respectively (Figure 4). This range of bacterial cell sizes were roughly equivalent to that of the wildtype on 2% agar.

**Fig. 4.**
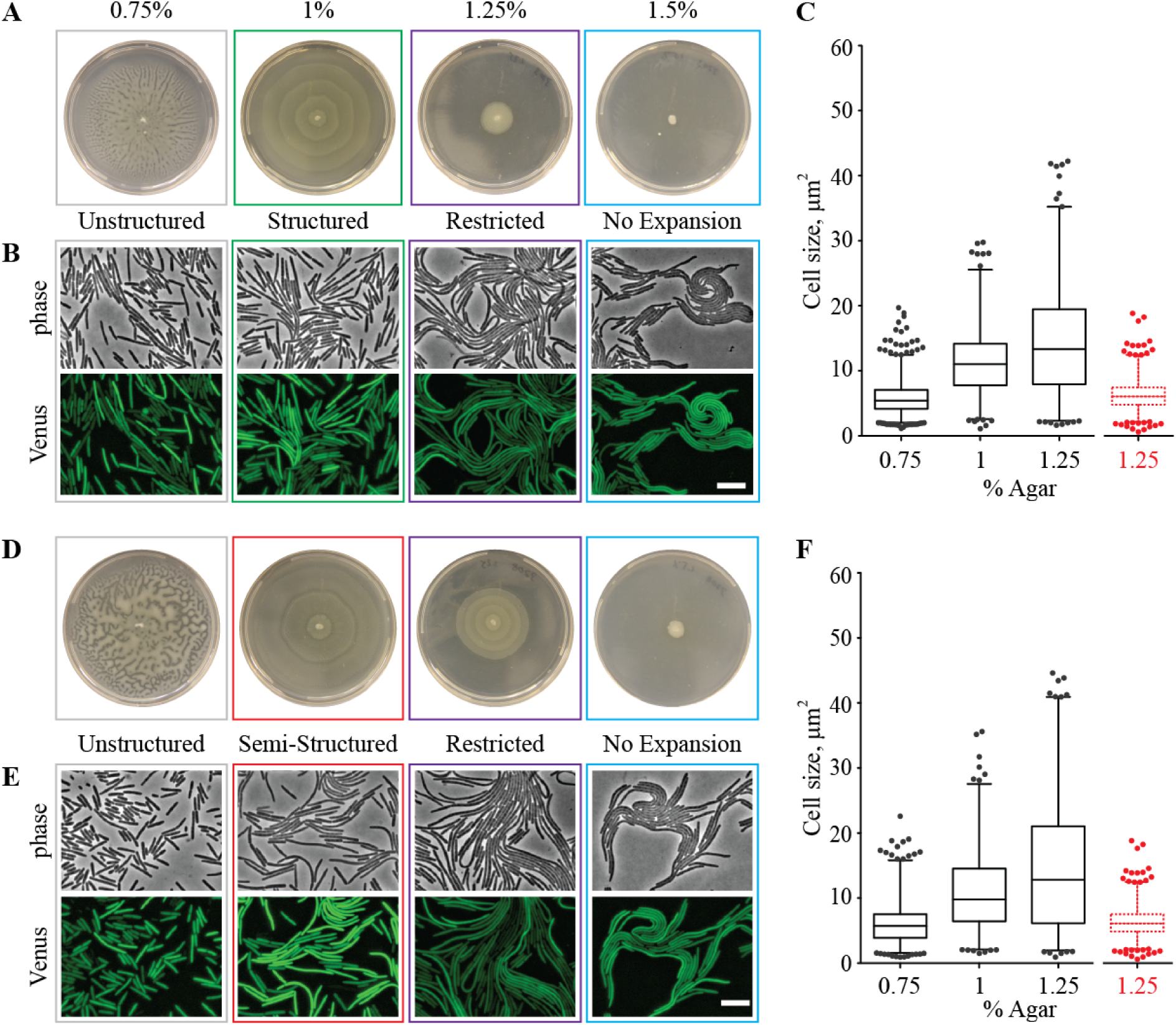
LPS-defective cells swarm and elongate more extensively than wildtype on lower-percentage agar. Representative images of strains KEL21 (A) and KEL22 (D), inoculated onto LB medium containing 0.75 to 4.0% agar and grown overnight at 37°C on a 10-cm diameter petri dish. Colony structure was categorized as unstructured, semi-structured, structured, and restricted as denoted earlier. Colonies with no swarm colony expansion are also indicated. Representative of three biological repeats. Representative images of strains KEL21 (B) and KEL22 (E) encoding a chromosomal Venus reporter for *fliA* expression. Cells were inoculated onto 1-mm thick LB medium pads containing 0.75 to 4.0% agar, grown at 37°C, and imaged using epifluorescence microscopy. Phase (top) and fluorescence (bottom). Fluorescence images were subjected to background subtraction. Scale bars = 10 µm. Representative of three biological repeats. Swarmer cell size on media containing varying concentrations of agar. KEL21 (C) and KEL22 (F) cells were co-mixed in a 10:1 ratio with corresponding cells carrying the *fliA* reporter strain and swarmed and analyzed and represented as described in Figure 2C. The numbers of measured cells for each agar percentage, sequentially from 0.75% to 1.25%, are as follows: (for KEL21) 2344, 795, and 823; (for KEL22) 1351, 757, and 796. Data for wildtype at 1.25% agar (1419 measured cells) is duplicated from Figure 2C and colored red for visual comparison.

The distribution of cell sizes scaled with agar concentration for populations of the mutant strains and were broader than the distributions of wild-type cell sizes at equivalent agar concentrations (Figures 1 and 4). For example, at 1% agar on which all populations could swarm, the range of sizes for KEL21 and KL22 was 28.7 µm^2^ and 34.1 µm^2^, respectively, as compared to 13.7 µm^2^ for the wildtype. At 1.5% agar, populations of each mutant strain were constrained to the site of inoculation with no visible terraces (Figure 4). In sum, single-cell morphology and the degree of cell-cell interactions corresponded to modifications to the visible structure of the swarm colony for both LPS-deficient cells and wild-type cells. By contrast, we found that a strain (KEL23), in which the flagella motor gene *motA* is disrupted, remained non-motile and failed to elongate into swarmer cells across all tested conditions (Supplemental Figure SF3). Taken together, we concluded that the LPS-deficient strains were not disrupted in elongation into swarmer cells and were instead specifically attenuated in population-wide swarm colony expansion.

### LPS-associated activity functions locally at individual cells to promote swarm colony expansion

A potential mechanism is that such LPS deficiencies inhibit a populations’ ability to acclimate to hardening surface conditions, such as agar concentrations of 1.5% and above. In such a case, LPS might function as a swarm-promoting public good in the extracellular matrix of the swarm colony to increase the swarming of neighboring cells. We therefore examined whether wild-type cells could rescue the swarm colony expansion for the LPS-deficient strains in trans, ostensibly through the sharing of the wild-type LPS. To employ differential measurement of swarm colony expansion (34), we engineered constitutive expression of carbenicillin resistance (Cb^R^) onto the chromosome of a wild-type population (wildtype::Cb^R^) and similarly engineered resistance to streptomycin (Sm^R^) onto the chromosome of the wildtype (wildtype::Sm^R^), KEL21 (KEL21::Sm^R^), and KEL22 (KEL22::Sm^R^) populations. We mixed the wildtype::Cb^R^ and the streptomycin-resistant wildtype or mutant strain populations at 1:1 or at 1:10 initial ratios (Figure 5). The mixtures were inoculated onto CM55 agar, LB 1.25% agar, or LB 1% agar and then grown for 20 hours at 37°C. A gridded sample of the swarm colony was then transferred onto non-swarm permissive selective media containing either 100 µg mL^-1^ carbenicillin or 25 µg mL^-1^ streptomycin to provide a coarse spatial measurement for swarm colony expansion.

**Fig. 5.**
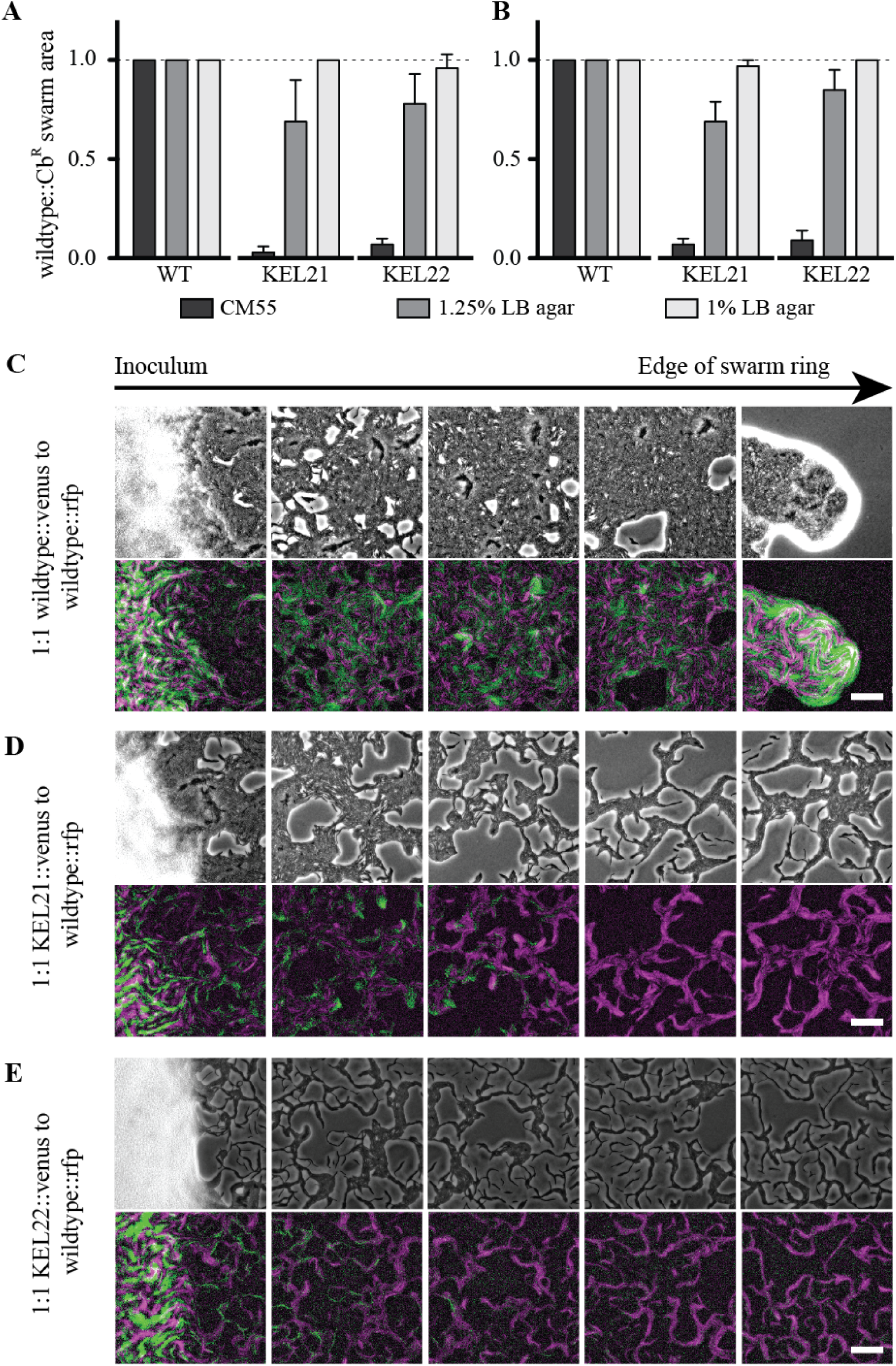
The swarm motility of LPS-deficient cells is not rescued by wildtype. Wildtype constitutively producing carbenicillin resistance (Cb^R^) were mixed at a 1:1 ratio (A) or a 1:10 ratio (B) with either the wild-type, KEL21, or KEL22 strains, each of which was constitutively producing streptomycin resistance (Sm^R^). Swarm area was calculated as the ratio of the area occupied by the Sm^R^ strain compared that occupied by the Cb^R^ strain. Error bars reflect standard deviation from the mean. Representative of three biological repeats. Wildtype constitutively producing mKate2 were mixed at 1:1 ratio with either wildtype(C), KEL21(D), or KEL22(E) derived strains carrying the chromosomal Venus reporter for *fliA* expression. Mixtures were swarmed on CM55 agar pads at 37 C. Phase images (top) and false-colored merge of fluorescence images (bottom) were taken once the first swarm ring formed. Images are taken scanning horizontally from the edge of the inoculum (white haze, far left image) to the leading front of the expanding swarm colony (far right image). mKate2-producing cells are false-colored magenta; Venus-producing cells are false-colored green. Fluorescence images were subjected to background subtraction. Scale bars, 50 µm. Representative of three biological repeats.

On CM55 agar, which is swarm-permissive for wildtype and swarm-restrictive for the LPS-defective strains, we found that populations of the KEL21 and KEL22 strains were restricted closer to the center while the wild-type population was at the leading edge regardless of initial ratio (Figure 5). On the LB agar conditions where all populations can swarm outwards when alone, the LPS-deficient strains were found near or at the leading edge along with the wildtype cells (Figure 5). We do not think that the restriction of LPS-deficient cells within a mixed swarm on CM55 agar was due to competition between wild-type and LPS-deficient populations for two reasons. First, altering the starting ratio to have 10 LPS-deficient cells to each wild-type cell did not impact the final result (Figure 5B). Second, LPS-deficient cells and wild-type cells co-swarmed completely on 1% and 1.25% agar (Figure 5). Instead, we interpreted these data to indicate that the presence of wild-type LPS structures did not rescue the swarm colony expansion of LPS-deficient populations, and therefore, LPS did not function as a secreted public good.

However, the observed lack of rescue could be caused by an inability of wild-type and LPS-deficient cells to closely associate. Therefore, we analyzed populations using epifluorescence microscopy to approach single-cell analysis. We mixed wild-type cells constitutively producing Red Fluorescent Protein (RFP) at a 1:1 ratio with either the wild-type, KEL21, or KEL22 strains that were engineered to encode the chromosomal Venus reporter for *fliA* expression. The mixed populations were inoculated onto CM55 agar and permitted to grow at 37°C. Upon emergence of the first swarm ring, swarm populations were imaged continuously in non-overlapping frames from the inoculation point to the leading edge of the actively migrating swarm (Figure 5). Cells swarmed equally together throughout the first swarm ring for the mixed wild-type strains, (Figure 5C). By contrast, cells producing Venus were restricted closer to the center than RFP-expressing wild-type cells for the mixed populations containing KEL21 or KEL22-derived strains (Figure 5D and 5E). Within rafts, swarming cells of all populations mixed, indicating that LPS-deficient cells could closely interact with wild-type cells. Similar results were obtained when wildtype-, KEL21-, and KEL22derived strains that were co-swarmed with wildtype at a 10:1 ratio (Supplemental Figure SF3).

Therefore, the defects associated with loss of wildtype LPS structures appeared to be constrained to individual cells. Wild-type cells could not rescue swarm motility of individual LPS-deficient cells even when physically aligned in an actively expanding swarm (Figure 5). We concluded that the LPS defects in strains KEL21 and KEL22 have severed the ability of individual cells to overcome environmental barriers to motility at 1.5% LB agar or higher; this might be by altering the interactions between each cell and the local rigid surface and/or neighboring cells. These LPS defects did not inhibit individual cell development into elongated swarmer cells and served instead to disrupt population-wide expansion by altering individual interactions with the immediate local environment.

## Discussion

We have found that swarming *P. mirabilis* responds to variations in the property of a surface by gradually increasing cell length and flexibility, cell-cell interactions, and overall swarm colony structure. Further, these micron-scale cellular changes correspond with centimeter-scale changes in the structure of the swarm colony. Increases in agar concentration beyond 1.5% were previously shown to reduce the speed as well as the time *P. mirabilis* populations spend expanding to form each terrace, thereby narrowing swarm colony terrace width (28, 39). More beyond this was not fully understood.

By examining populations grown across a broad range of surface conditions at both macroscopic and microscopic scales, we have uncovered several important aspects. First, surface-grown *P. mirabilis* cells elongate across a gradient of sizes and form more extensive cell-cell contacts in a dynamic response to increasing agar concentration and likely surface hardness (Figure 6A). Second, changes in swarm colony structure and terrace size reflect these microscopic changes; indeed, the degree of cell elongation across a population correlated with the visible structure of the swarm colony. Third, LPS functions locally to help individual cells to overcome environmental barriers to motility. These motility barriers may be related to surface tension, agar wetness, or osmotic pressure (3, 22, 23, 27). Loss of LPS-associated structures restricts swarm colony expansion to occurring in lower-density agar conditions and leads to enhancement in cell elongation (Figure 6B). We propose a coarse model in which swarmer cell morphology and cell-cell interactions scale in response to barriers to motility (Figure 6). The visible changes in swarm colony structure likely arise from the micrometer-scale phenotypes.

**Fig. 6.**
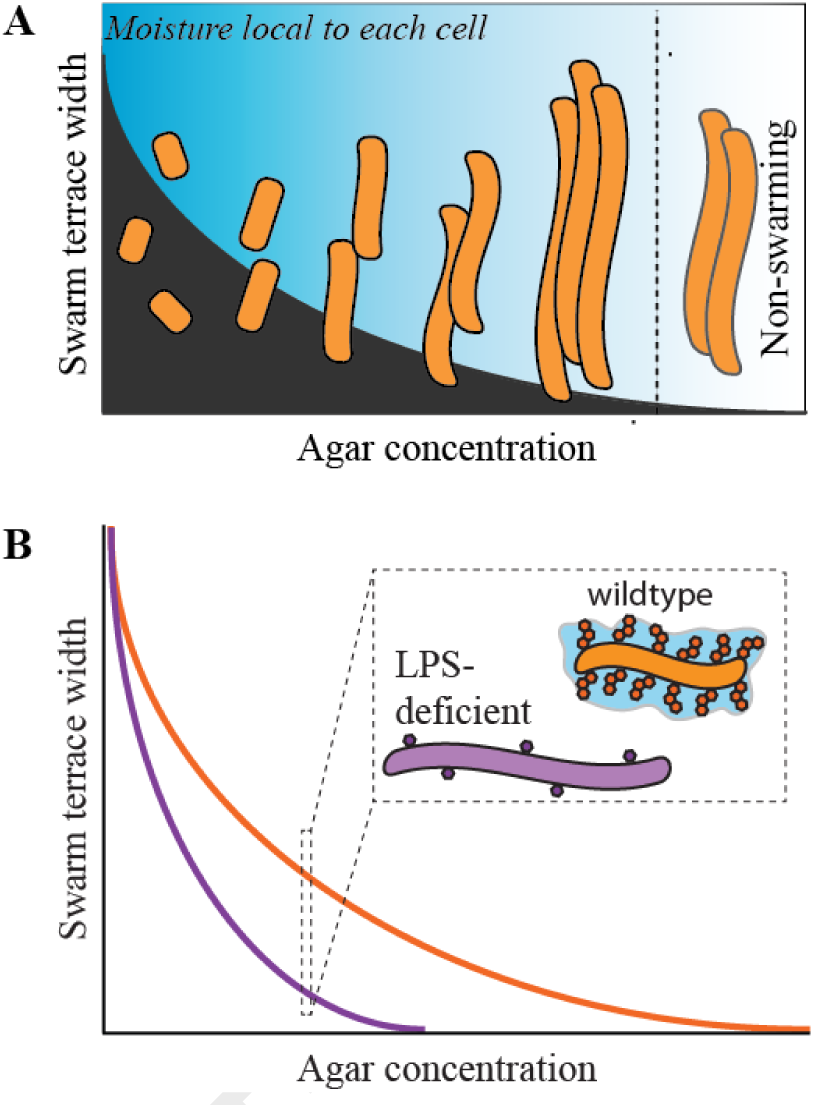
Model for how *P. mirabilis* cell morphology and cell to cell interactions respond to surfaces. A. We propose that *P. mirabilis* has a gradient of responses to agar concentration: gross modifications to cell morphology, cell-cell alignment, and macroscale colony structures. We hypothesize that agar-associated barriers to motility may be related to local moisture levels and/or surface hardness. B. Summary graphic depicting that LPS-deficient cells elongate more extensively than wildtype cells on a given agar medium. Coordinately, populations of LPS-deficient cells form swarm colonies with more restricted terrace structures than wild-type populations and engage in swarm colony expansion across a narrower range of agar conditions.

The relationship between single-cell morphology, cell behaviors, and population-wide swarm colony expansion remain unclear. Here we have shown that increased cell length and degree of cell-cell association strongly correlated with centimeter-scale swarm colony architecture for both wildtype and LPS-deficient populations. Indeed, the emergence of a terraced swarm colony structure of wild-type populations on 1.25% agar correlated with more frequent and extensive associations between cells at the micron scale. Further remains to be done to establish how the movements of individual cells drive the emergence of differential macroscopic population structures. Given that the degree of cell elongation across a population is predictive of swarm colony structure, one wonders whether additional, currently uncharacterized factors also influence the emergence of swarm colony expansion.

We further propose that novel and yet uncharacterized mechanisms exist to tune swarmer cell elongation and promote multicellular rafting based on integrated environmental cues. The *motA*-deficient strain failed to elongate, and flagellar activity is linked to swarmer cell development (40). Therefore, such unidentified mechanisms presumably function downstream of a theorized flagella-mediated surface-sensing switch. One critical insight derived from this research— that swarmer cell elongation can be uncoupled from swarm colony expansion through increasing agar concentration or altering LPS structure— provides a first step for elucidating such swarm-promoting factors.

In addition, several bacterial species modify their local environment and cooperative behaviors to promote swarm colony expansion. For example, surface-motile cells appear to migrate along shared tracks comprised of furrows in the medium lined with extracellular material in *Pseudomonas aeruginosa* and *M. xanthus* (41–43). In *P. mirabilis*, swarm colonies can produce extracellular slime trails that appear to spatially coordinate swarming cells to promote swarm colony expansion (44). In addition, LPS has been proposed to be shed by cells, potentially sheared off by flagella movement, to act as a shared-good surfactant (31).

By contrast, the results here indicate that LPS functions locally to individual *P. mirabilis* cells. LPS appears to act on individual cell motility, likely independent of the creation of slime tracks or burrows under various swarm conditions. Co-incident swarming with wild-type cells, presumably capable of creating slime tracks or burrows, did not rescue the colony structure and cell motility of LPS-deficient mutant populations. Yet cells of the mutant strains could elongate and align into motile rafts with wild-type cells. The two populations equally swarmed outwards on low-agar medium permissive to the swarm expansion of mutant strains. Even though wildtype did not rescue the mutant strain for swarming *in trans* on higher agar conditions, active exclusion by the wildtype was also not apparent. We cannot formally rule out the possibility that LPS might interact with other secreted factors; this remains to be investigated. We propose that LPS surface molecules could possibly reduce environmental barriers to swarming by locally altering each individual cell’s interactions with the environment, perhaps increasing the ability of an individual cell to draw moisture from the medium (Figure 6).

In a broader context, the ability of bacteria to tune cell morphology and behavior might contribute to survival as cells acclimate to changing environments. *P. mirabilis* is an opportunistic pathogen and commonly found resident of human and animal intestines. Bacteria encounter variable chemical, immunological, and mechanical environments within a host organism (45). In the specific context of *P. mirabilis*, swarmer cells might be important for invasion of a catheterized bladder; within the bladder, formation of short cells and biofilms might be more beneficial for long-term infections. Indeed, increasingly elongated cells produce more flagella and form a greater number of cell-cell contacts, allowing for rapid migration, than shorter cells. However, increased cell length in and of itself may not be a positively selected adaptation, but rather a secondary or neutral effect. For example, increased cell elongation may be a secondary effect of increased flagella production, which in itself provides a clear motility-associated advantage for greater motion per cell but can also cause challenges within a host as flagella and LPS structures are known to activate host immune responses. The balance and tuning of flagella, LPS, cell shape, and population motility, especially over the course of invasion and infection, remains unresolved. Further investigations at the interface of outer membrane structure, motility, and cell morphology could lead the field toward more clinically relevant insights into bacterial behavior and potential identification of therapeutic targets.

## Methods

### Media

Liquid cultures were grown in Lennox lysogeny broth (LB). Colonies were grown using LB containing Bacto Agar, from 0.75% to 4% agar, or using CM55 agar (Oxoid, Hampshire, UK) (1.5% agar) for motility assays as indicated. LSW- agar (46) was used for plating non-motile colonies. For swarm assays, overnight cultures were normalized to an optical density at 600 nm of 1.0, and culture was then inoculated with an inoculation needle onto the medium before growth at 37°C.. Antibiotics used were: 100 µg mL^-1^ tetracycline, 50 µg mL^-1^ chloramphenicol, 100 µg mL^-1^ carbenicillin, and 35 µg mL^-1^ kanamycin. Dyes were added to swarm plates where indicated: 40 µg mL^-1^ Congo Red and 20 µg mL^-1^ Coomassie Blue.

### Strains and plasmids

Wild-type strain *P. mirabilis* strain BB2000 (46) and wildtype producing RFP (47) were previously described. The mutant strains KEL21 (BB2000 with a Tn-Cm^R^ disruption of BB2000_3203) and KEL22 (with a Tn-Cm^R^ disruption of BB2000_3208) are first described in this study. Each strain was generated through mutagenesis of strain BB2000 with mini-Tn5 carrying a chloramphenicol-resistance marker within the transposable element and a carbenicillin-resistance marker on the vector backbone as previously described (46). The locations of transposon insertions in strains KEL21 and KEL22 were determined through inverse polymerase chain reaction (PCR) following the previously described protocol for P. mirabilis (34). Briefly, genomic DNA was extracted from each strain through phenol/chloroform extraction. Genomic DNA was digested with HhaI, and the resultant fractions ligated with T4 ligase overnight. Transposon-specific primers coupled with random primers were used to PCR amplify a region flanking the 5’ and separately 3’ regions of the insertion site. The resultant PCR products were purified, subjected to Sanger sequencing with transposon-specific primers by Genewiz, Inc. (Cambridge, MA), and mapped to the BB2000 genomic sequence. Protein sequences were analyzed using HMMER (phmmer) (**?**) and the UniProtKB database (48).

Strain construction was performed as described in (49). For all strains, expression plasmids were introduced into *P. mirabilis* via conjugation with *E. coli* SM10 lambda pir (46). Resultant strains were confirmed by Polymerase Chain Reaction (PCR) of the targeted region. For construction of *fliA* reporter strain, a gBlock (Integrated DNA Technologies, Skokie, IL) encoding the last 500 base-pairs (bp) *fliA* (*P. mirabilis* BB2000, accession number CP004022:nt 1856328… 1856828), an RBS (aggagg), a modified variant of Venus fluorescent protein (50), and 500 bp downstream *fliA* (*P. mirabilis* BB2000, accession number CP004022:nt 1855828… 1856328) was inserted into pKNG101 /citeKaniga1991 at the ApaI and XbaI sites. For construction of the streptomycin-resistant strains, a constitutive mKate2 expression construct flanked by regions homologous to *rluA* (BB2000_2998) and BB2000_2999 was generated using a gBlock (Integrated DNA Technologies, Skokie, IL) and inserted into pKNG101 at the ApaI and SpeI sites. For generation of the carbenicillin-resistant strains, a constitutive *gfpmut2* expression construct flanked by regions homologous to *rluA* and BB2000_2999 was generated by SOE PCR (51) and inserted into pRE107 (52) at the KpnI and XmaI sites. Resultant strains were maintained as merodiploids. All plasmids were confirmed by Sanger Sequencing (Genewiz, South Plainfield, NJ).

### Antibiotics susceptibility assay

Cultures were top-spread on LSW- medium and allowed to sit at room temperature until surface appeared dry (approximately 2 hours). Then, 6-mm sterile filter disks were placed onto plates and soaked with 10 µL of polymyxin B (Sigma Aldrich, St Louis, MO) solution. A water-alone control was included. Once filter disks dried (approximately 2 hours), plates were incubated at 37°C overnight and imaged. The minimal inhibitory concentrations were determined based on the concentration of polymyxin B that produced a clearing in the bacterial lawn.

### LPS extraction and analysis

Cells were grown on overnight at 37°C on CM55 agar, and the resultant swarms harvested with LB broth as previously described (32). LPS was extracted from cells using an LPS Extraction Kit according to manufacturer’s instructions (iNtRON Biotechnology Inc, Sangdaewon Seongnam, Gyeonggi, Korea). Extracts were resuspended in 10mM Tris, pH 8.0 buffer and run on a 12% SDS-PAGE gel. Gels were stained using silver stain (36).

### Microscopy

Microscopy was performed as previously described (53). Briefly, agar pads were inoculated from overnight stationary cultures and incubated at 37°C in a modified humidity chamber. Pads were imaged using a Leica DM5500B (Leica Microsystems, Buffalo Grove, IL) and a CoolSnap HQ2 cooled CCD camera (Photometrics, Tuscon, AZ). MetaMorph version 7.8.0.0 (Molecular Devices, Sunnyvale, CA) was used for image acquisition. Images were analyzed using FIJI (54). Where indicated, images were subjected to background subtraction equally across the entire image. Venus (max excitation 515 nm; max emission 528 nm) was visualized using a GFP ET filter cube (excitation 470/40 nm; emission 525/50 nm) (Leica Microsystems, Buffalo Grove, IL). Corresponding movies can be found on the K. A. Gibbs’ Research Group’s YouTube channel.

### LPS extraction and analysis

Cells were grown on overnight at 37°C on CM55 agar, and the resultant swarms harvested with LB broth as previously described (32). LPS was extracted from cells using an LPS Extraction Kit according to manufacturer’s instructions (iNtRON Biotechnology Inc, Sangdaewon Seongnam, Gyeonggi, Korea). Extracts were resuspended in 10mM Tris, pH 8.0 buffer and run on a 12% SDS-PAGE gel. Gels were stained using silver stain (36).

### Quantifying swarmer cell size

Mixed swarms of *fliA* reporter encoding and non-encoding populations were mixed at indicated ratios and allowed to swarm on LB pads containing the indicated amount of agar at 37°C until the first swarm ring was visible by eye. Swarms were imaged using phase and epifluorescence microscopy. We utilized MATLAB and Image Processing Toolbox Release 2018a (The MathWorks, Inc., Natick, MA) to assemble a pipeline for analyzing swarmer cell size from microscopy images. The source for basic structure of the segmentation code, specifically the section that thresholds the image, was based on http://blog.pedro.si/2014/04/basic-cell-segmentation-in-matlab.html (55). The source for utilizing other algorithms used in code as well as the assistance in implementing was was The Mathworks, Inc. Image Processing Toolbox: Reference (r2018a), which can be found at http://www.mathworks.com/help/pdf_doc/images/images_ref.pdf. Data was filtered by eye to only include intact, individual cells in the analysis.

### Mixed swarm assays

Overnight cultures were normalized to an optical density at 600 nm of 1. For 1:1 and 1:10 mixed populations, samples were prepared at the appropriate ratio from the normalized cultures. Topological mapping was performed as described as detailed in (34). Mixed populations were inoculated onto CM55 agar, 1.25% LB agar, or 1% LB agar and then allowed to swarm at 37°C overnight. Once the swarm reached the plate’s edge, a 48-pronged replicator device was used to sample regions of the swarm and then replica-plated onto two independent plates: one containing streptomycin and the other containing carbenicillin. These antibiotic-containing plates were grown overnight at 37°C and imaged with a Canon EOS 60D camera. The number of spots corresponding to each of the 48 prongs was counted. The reported ratio value is the number of streptomycin growth spots over the number of carbenicillin growth spots. For microscopy, mixed cultures were allowed to swarm on CM55 agar plates until the first swarm ring was visible by eye. Plates were then imaged at 40X magnification as described above. Images were taken continuously in non-overlapping frames from the inoculation point to the leading edge of the actively migrating swarm.

## ACKNOWLEDGEMENTS

We would like to thank Drs. Enrique Balleza and Philippe Cluzel for the gift of the Venus construct and John Welsh for assistance in constructing the wild-type derived *fliA* reporter strain and strain KEL23. We would like to thank members of the Gibbs, D’Souza, Gaudet, and Losick research groups for thoughtful discussion and feedback. This research was funded by a Smith Family Graduate Fellowship in Science and Engineering, the David and Lucile Packard Foundation, the George W. Merck Fund, and Harvard University.

## AUTHOR CONTRIBUTIONS

KL and KAG conceived and designed the study and wrote the manuscript. JA compiled the workflow used to measure cell area. JZ conducted a subset of swarm assays. KL conducted all other experiments.

## Supplementary Note 1: Supplemental methods

### Strains and media

Liquid cultures were grown in Lennox lysogeny broth (LB). Colonies were grown using LB containing Bacto Agar, from 0.3 to 4% agar or using CM55 agar for motility assays as indicated. LSW- agar (46) was used for plating non-motile colonies. For swarm assays, overnight cultures were normalized to an optical density at 600nm of 1.0, and the culture was inoculated with a needle onto the medium and grown at 37°C. Antibiotics used were: 100 µg mL^-1^ tetracycline, 50 µg mL^-1^ chloramphenicol, 100 µg mL^-1^ carbenicillin, and 35 µg mL^-1^ kanamycin. Dyes were added to swarm plates where indicated: µg mL^-1^ Congo Red and 20 µg mL^-1^ Coomassie Blue.

For construction of the *motA* disruption in strain KEL23, a gBlock (Integrated DNA Technologies, Skokie, IL) encoding the last 500 bp of *flhC* (*P. mirabilis* BB2000, accession number CP004022:nt complement 1908501… 1909001), a ribosomal binding site (aggagg), the gene encoding a modified Cyan Fluorescent Protein (50), and the 500 bp downstream of *flhC* (*P. mirabilis BB2000*, accession number CP004022:nt complement 1908001… 1908501) was inserted into pKNG101 at the ApaI and XbaI sites. The resultant strains were maintained as merodiploids and *in vivo* fluorescence was evaluated with epifluorescence microscopy. This method was otherwise performed as described in the main text. Media conditions were used as described in the main text.

### Mixed swarm microscopy assay

Overnight cultures were normalized to an optical density at 600 nm of 1. For 1:10 mixed populations, samples were prepared at the appropriate ratio from the normalized cultures, inoculated onto CM55 agar, and allowed to swarm at 37°C until the first swarm ring was visible by eye. Plates were then imaged at 40X magnification. Images were taken continuously in non-overlapping frames from the inoculation point to the leading edge of the actively migrating swarm. Microscopy was performed as previously described (53). Briefly, agar pads were inoculated from overnight stationary cultures and incubated at 37°C in a modified humidity chamber. Pads were imaged using a Leica DM5500B (Leica Microsystems, Buffalo Grove, IL) and a CoolSnap HQ2 cooled CCD camera (Photometrics, Tuscon, AZ). MetaMorph version 7.8.0.0 (Molecular Devices, Sunnyvale, CA) was used for image acquisition. Images were analyzed using FIJI (54). Where indicated, images were subjected to background subtraction equally across the entire image. Venus (max excitation 515 nm; max emission 528 nm) was visualized using a GFP ET filter cube (excitation 470/40 nm; emission 525/50 nm) (Leica Microsystems, Buffalo Grove, IL).

## Supplementary Note 2: Supplemental figures

**Figure SF1.**
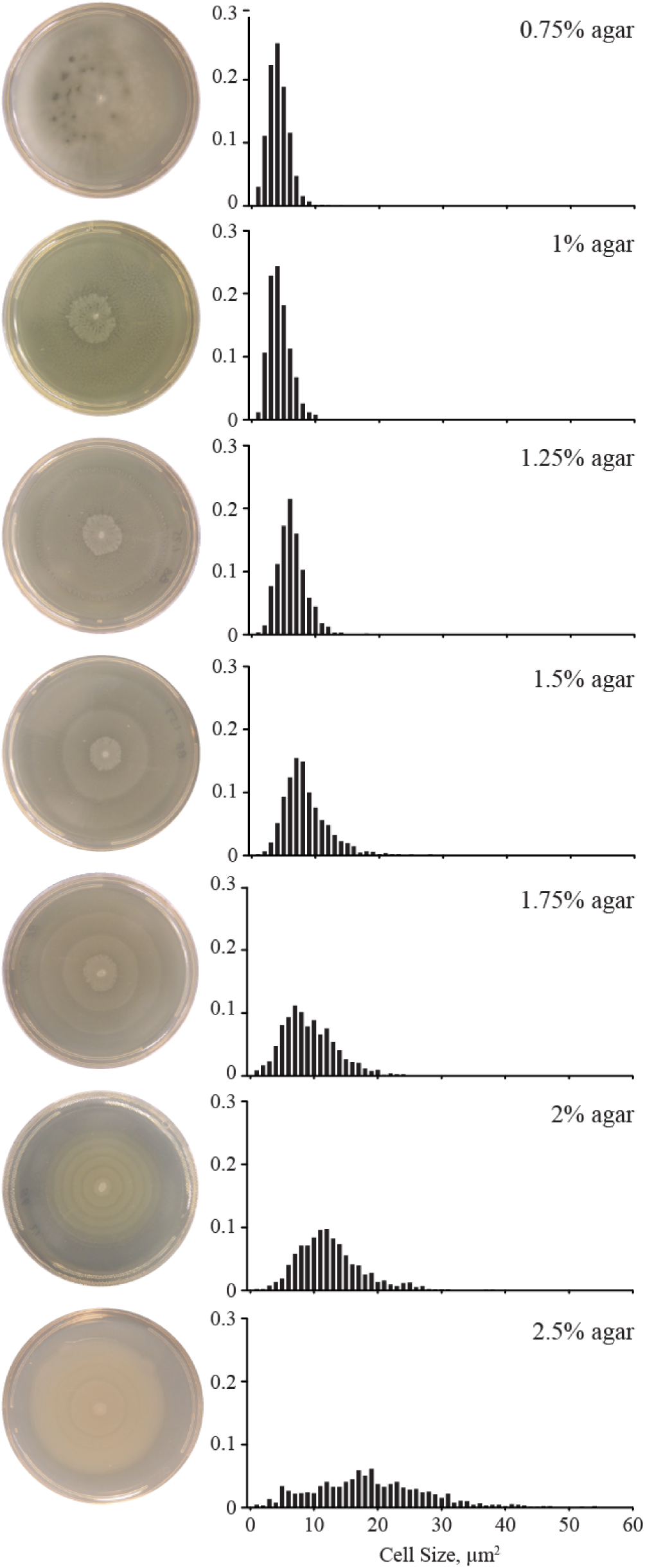
Cell size and swarm colony structure shifts as agar concentration Increases. A. Representative images of wild-type *P. mirabilis* that were inoculated onto LB medium containing the indicated agar concentration and then grown overnight at 37°C on a 10-cm diameter petridish. The width of individual terraces decreased as agar concentrations increased. Terraces were not readily apparent at 0.75% agar. B. The area of individual swarmer cells was calculated on different agar surfaces. The plotted graphs are histograms of the datasets. The data for Y-axis denotes the fraction of cells within each poplation. Bin size, 1 μm^2^. The data for 0.75%, 1%, 1.25%, 1.5%, and 2.5% are presented in the main text.

**Fig. SF2.**
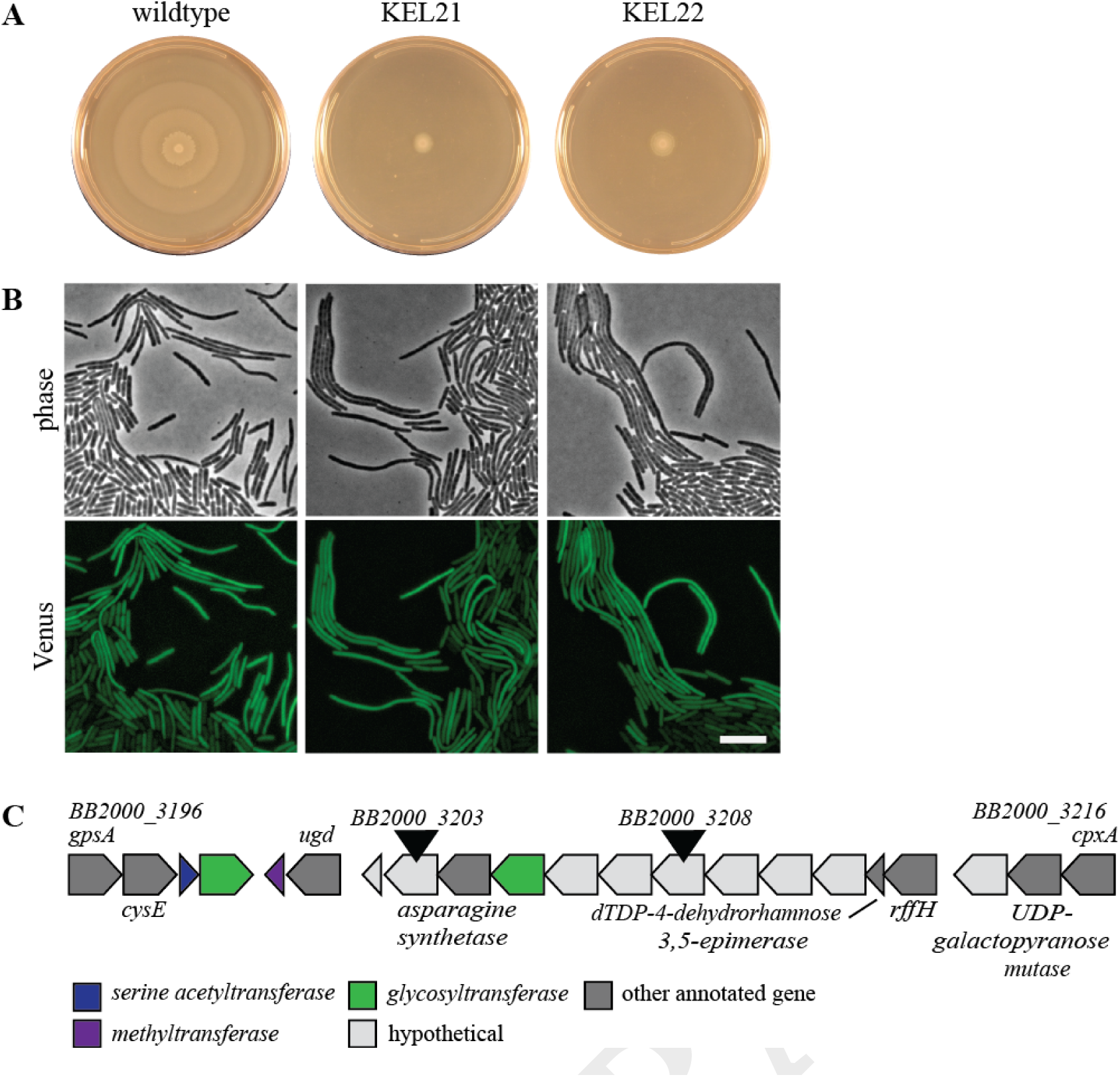
Disruption of LPS biosynthesis genes uncouples swarmer cell elongation and swarm colony expansion. A. Wildtype and two mutant strains (KEL21and KEL22) inoculated onto CM55 swarm medium and grown overnight at 37 C. The wild-type population formed bullseye-patterned swarm colonies, while the mutant strains formed colonies constrained to the site of inoculation. Representative of three biological repeats. B. Representative images (from left to right) of wildtype, and strains KEL21 and KEL22; each strain encodes a chromosomal Venus reporter for *fliA* expression. Cells were inoculated onto 1-mm thick CM55 agar pads, grown at 37°C, and imaged using epifluorescence microscopy. Phase (top) and in fluorescence (bottom). Background subtraction was performed across the entire fluorescence images. Scale bars, 10 µm. Representative of three biological repeats. C. The gene cluster disrupted in KEL21 and KEL22 is predicted to be needed for lipopolysaccharide (LPS) biosynthesis (35) and varies between strains. Black arrows mark the genes independently disrupted in strains KEL21 and KEL22, respectively. Other gene names found below. Figure is not drawn to scale.

**Fig. SF3.**
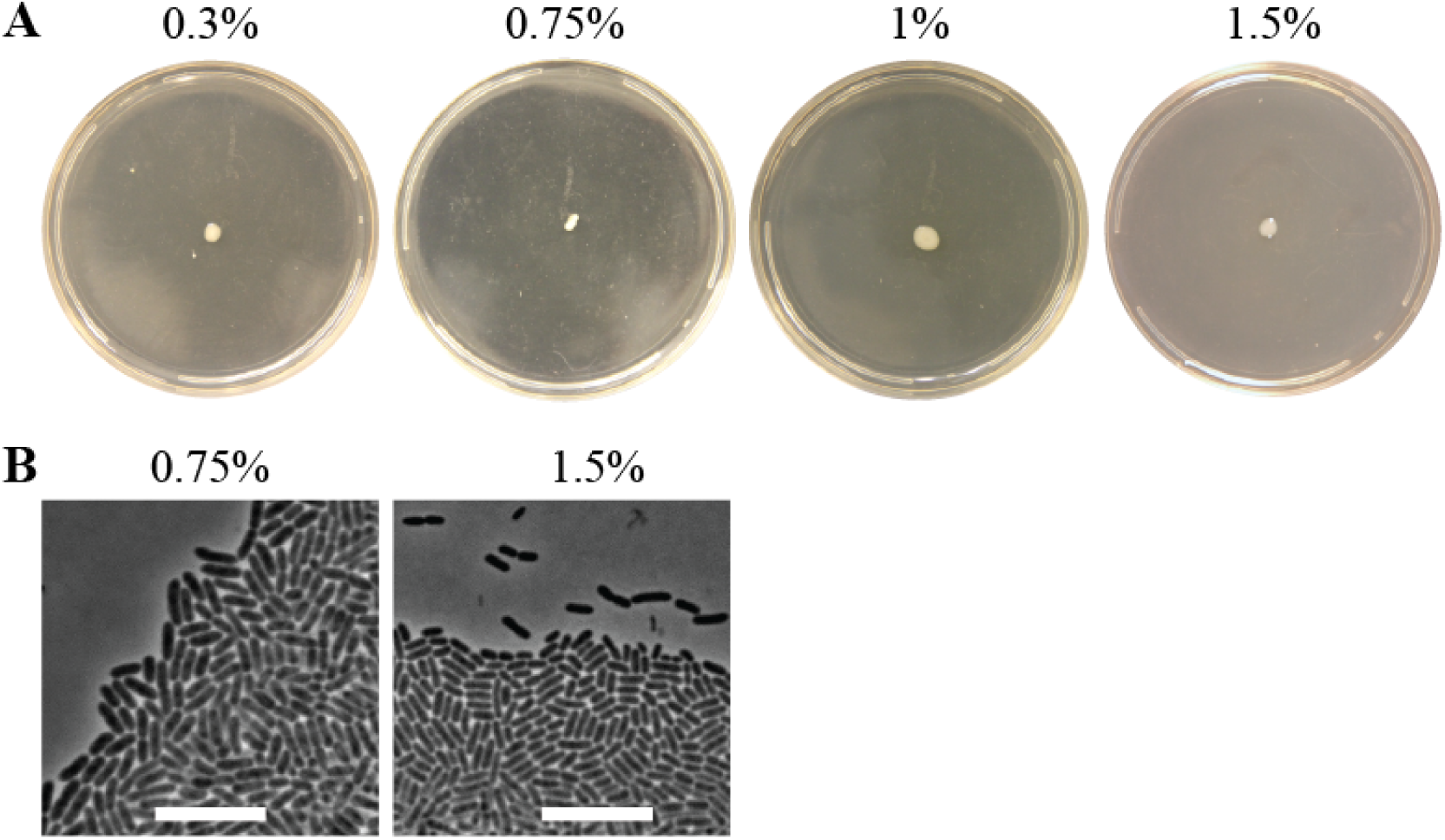
Disruption of the *motA* gene inhibits swarmer cell elongation and motility. A. Strain KEL23, which encodes a disruption of *motA* expression, was cultured in LB broth, and then swarmed on LB agar containing indicated concentrations of agar. Representative of three biological repeats. B. Representative image of the colony edge for strain KEL23 after 8 hours of incubation at 37 C on the indicated agar medium. Representative of three biological repeats. Scale bars, 10 µm.

**Fig. SF4.**
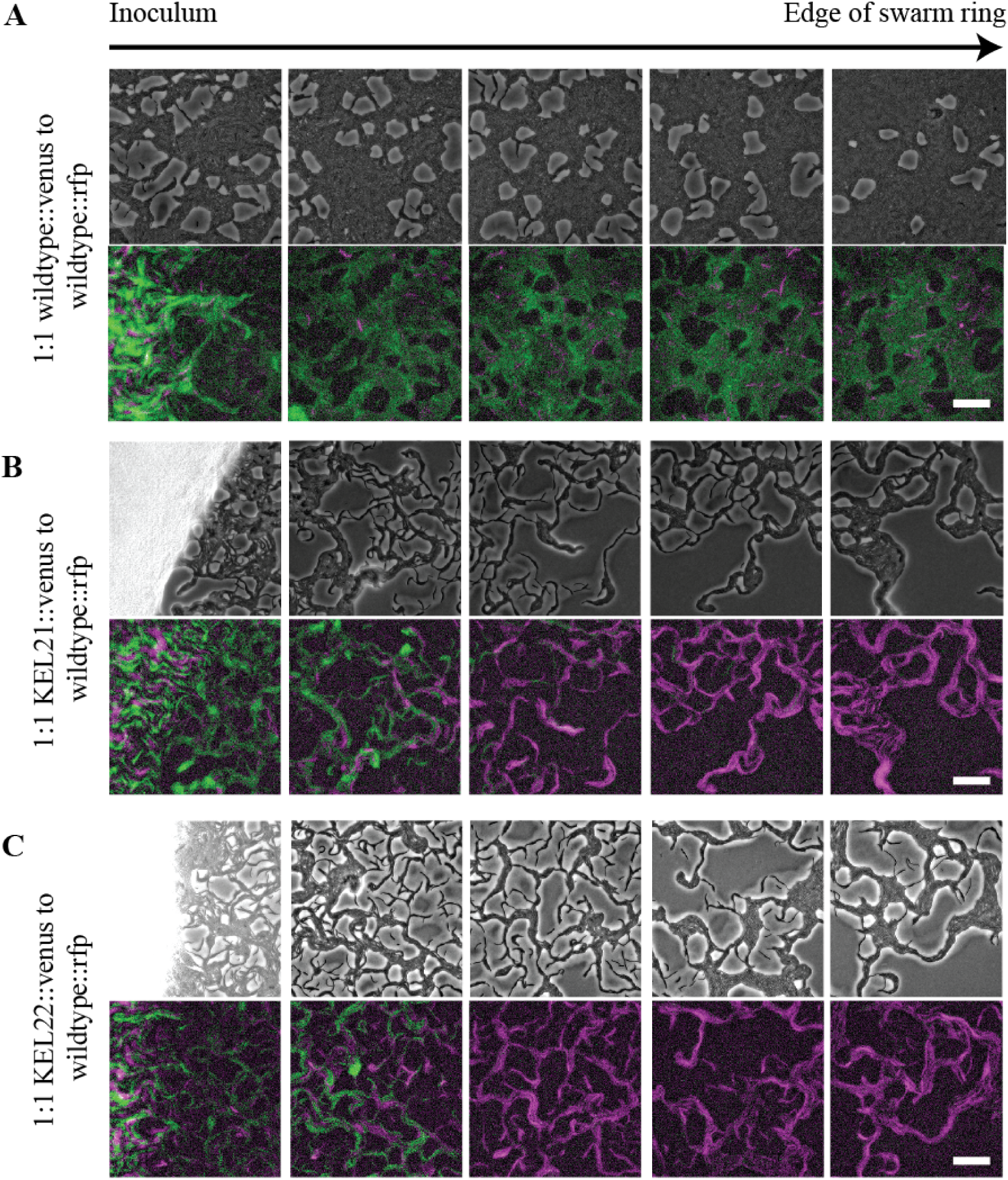
LPS-deficient strains remain proximal to inoculation point when mixed with wild-type populations. Wildtype constitutively producing mKate2 was mixed at 1:10 ratio with either the wildtype-(A), KEL21- (B), or KEL22- (C) derived strains carrying the chromosomal Venus reporter for *fliA* expression. mKate2-producing cells are false-colored in magenta; Venus-producing cells are false-colored in green. Phase images (top) and false-colored merge of fluorescence images (bottom) were taken once the first swarm ring formed, scanning horizontally from the edge of the inoculum (white haze, far left image) to the leading front of the expanding swarm colony (far right image). Fluorescence images were subjected to background subtraction. Scale bars, 50 µm. Representative of three independent experiments per condition.

